# Epileptogenesis dynamics driven by peritumoral circuit rewiring in gangliogliomas

**DOI:** 10.64898/2026.08.04.742675

**Authors:** Silvia Cases-Cunillera, Louise Deboeuf, Alesya Evstratova, Joshua Reid, Belén Díaz-Fernández, Jan Pyrzowski, Juan Nieto, Kirill Smirnov, Demiana Fanous, Johan Pallud, Thomas Blauwblomme, Rainer Surges, Michel Le Van Quyen, Elena Dossi, Albert J. Becker, Gilles Huberfeld

## Abstract

Gangliogliomas (GGs) are emblematic low-grade epilepsy-associated tumors, yet the developmental mechanisms underlying their epileptogenicity remain unclear. Here, we investigated how tumor-network interactions evolve across postnatal maturation using an in-utero electroporated BRAF^V600E^-driven mouse model combining multiscale electrophysiology with histology, single-nucleus RNA sequencing, and complementary analyses in human GG tissue. We show that GGs induce early and evolutive modification of cortical organization and glioneuronal architecture. Despite glioneuronal preservation, seizure initiation shifted from distal cortical regions at postnatal stages to tumor-adjacent areas in adult networks. In both mouse and human, GG slices exhibited seizure activity localized to the peritumoral cortex. At the cellular level, neurons exhibited a developmental arrest of intrinsic electrophysiological maturation from postnatal to adult stages. Transcriptomic profiling identified stage-specific neuronal remodeling, with early alterations in inhibitory neurons and later changes affecting excitatory populations. Such developmental spatial seizure dynamics were associated with a pharmacological shift as NKCC1 inhibition with bumetanide selectively reduced seizure-like activity in neonatal but not mature tumor networks, indicating a restricted window of chloride-dependent epileptogenesis relevant to GABAergic maturation. Together, our results demonstrate that GG-associated epileptogenesis arises from developmentally regulated tumor-network interactions, highlighting distinct cellular and molecular mechanisms across maturation and revealing potential age-specific therapeutic targets.

## Introduction

Gangliogliomas (GGs) are among the most common low-grade epilepsy-associated tumors (LEATs), characterized by a mixed glioneuronal composition and a strong association with early-onset, frequently drug-resistant focal epilepsy, particularly in children and young adults (Slegers and Blumcke, 2020; Thom et al., 2012). Although GGs are generally considered benign from an oncological perspective, their clinical relevance is largely driven by their epileptogenic potential. Notably, seizure freedom is not universally achieved following gross total resection, with a substantial proportion of patients continuing to experience postoperative seizures (Aronica et al., 2001; Blumcke et al., 2014; Englot et al., 2012), suggesting that the epileptogenic zone can extend beyond the tumor itself and may involve peritumoral cortical networks. Indeed, structural and functional abnormalities have been identified in the peritumoral cortex of GGs, including altered neuronal excitability and impaired inhibitory signaling (Aronica et al., 2007b).

GGs exhibit several features suggestive of a developmental origin, including their early clinical presentation, association with chronic epilepsy, and the presence of dysplastic neuronal elements and progenitor-like cell populations (Blümcke and Wiestler, 2002; Regal et al., 2023). In line with this developmental context, recent experimental studies suggest that GG-associated oncogenic signaling can actively perturb neuronal differentiation, synaptic integration, and circuit function (Koh et al., 2018; Müller et al., 2024), thereby contributing to epileptogenesis beyond the effects of a purely structural or mechanical lesion. These findings support a model in which tumor cells dynamically interact with the surrounding neural network, thereby influencing its maturation and excitability.

Despite these advances, the temporal and spatial dynamics by which GGs shape neuronal circuits and generate epileptogenic networks remain poorly understood, which is key to both identifying the dynamics of epileptogenesis processes and interfering with brain maturation. In particular, it is unclear how tumor– network interactions evolve across development, how seizure initiation relates to tumor versus peritumoral regions, and whether distinct mechanisms underlie early versus later disease stages. Here, we address these questions using a developmental mouse model of BRAF^V600E^-driven GG (Cases-Cunillera et al., 2022) combined with multiscale electrophysiology and single-nucleus transcriptomics and complemented by recordings from human GG tissue, to identify stage-specific mechanisms that may uncover novel therapeutic targets for seizure activity.

## Results

### Early and sustained glioneuronal disorganization and tumor-cell-specific loss of neuronal identity

Given the early clinical presentation of GGs and their glioneuronal composition, we characterized key tumor–microenvironment interactions already established during early brain development. To examine how GG cells shape their microenvironment, we utilized a previously established mouse model based on in-utero electroporation (IUE) (Cases-Cunillera et al., 2022). Tumors were analyzed at two developmental stages, postnatal day 5 (p5, GG^p5^) and young adulthood p20 (GG^p20^), and compared with age-matched IUE controls expressing fluorescent protein only (Ctrl^p5^, Ctrl^p20^; **Fig. 1A**). Immunostaining for GFAP revealed abundant reactive astrocytes within the tumor core and extending into adjacent tissue at both time points (**Fig. 1B**). To quantify astrocytic reactivity, images encompassing the IUE regions were overlaid with a grid of equally sized square regions (0.04 mm² each), and the percentage of GFAP-positive area was determined for each square individually (n=6-10 slices, from 4-6 mice). Control brains exhibited generally low GFAP immunoreactivity, with most ROIs containing only small GFAP-positive fractions. In contrast, BRAFV600E/pAkt tumors showed markedly increased GFAP staining, with some ROIs reaching up to ∼80–90% GFAP-positive area. GFAP enrichment was most prominent in ROIs located close to the IUE region and tended to decline with increasing distance from the IUE site (**Fig. 1C**). Elevated GFAP immunoreactivity was observed at both p5 and p20, suggesting that tumor-associated astrocytic reactivity is established early after birth and persists during postnatal maturation.

**Figure 1.**
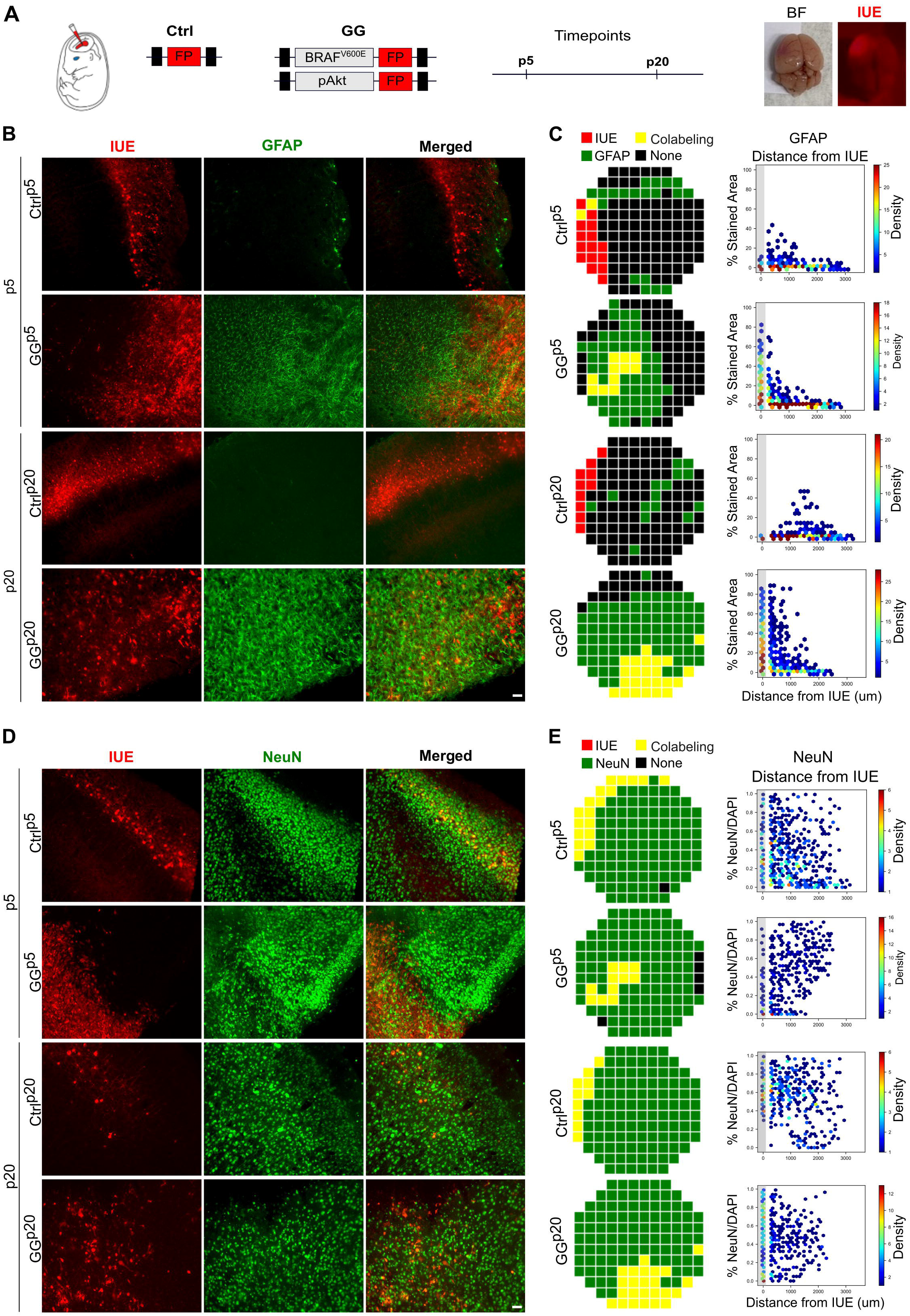
Representative immunofluorescence of glioneuronal markers in Ctrl and GG tissue. (**A**) Experimental design of in-utero electroporation (IUE), plasmids used to generate Ctrl and GG models, and postnatal timepoints analyzed. Representative brightfield and fluorescence images of whole IUE-electroporated brains are shown. (**B**) Immunofluorescence for GFAP (green) and the IUE fluorescent reporter (red) in Ctrl and GG cortical regions at p5 and p20. Scale bar, 50μm. (**C**) Grid-based spatial maps showing the distribution of IUE+ cells (red), GFAP+ astrocytes (green), and colabeled cells (yellow) in control (Ctrl) and GG brains at p5 and p20. Corresponding scatter plots depict the relationship between distance from IUE+ regions and percentage of GFAP-stained area, indicating increased astrocyte presence near tumor regions, particularly at p20. (**D**) Immunofluorescence for NeuN (green) and the IUE fluorescent reporter (red) from Ctrl and GG cortical tissue at p5 and p20. Scale bar, 50μm. (**E**) Spatial maps of IUE+ cells (red), NeuN+ neurons (green), and colabeled cells (yellow) across conditions. Scatter plots show NeuN/DAPI ratios as a function of distance from IUE+ regions. n=6-10 slices, from 4-6 mice.

To assess the neuronal compartment, NeuN immunostaining was performed and confirmed the presence of neurons within the tumor milieu at both developmental stages (**Fig. 1D**). Neuronal abundance was quantified by determining the proportion of NeuN-positive cells relative to all DAPI-positive nuclei within individual 0.04 mm² regions distributed across the analyzed sections. This analysis revealed no significant differences between control and tumor tissues, indicating that neuronal abundance was not significantly altered in BRAF^V600E^/pAkt tumors (n = 6 slices from 4 mice; **Fig. 1E**). Overall, these findings indicate that GG cells shape the glioneuronal landscape from early postnatal stages onward. This remodeling is characterized by a robust and sustained astrocytic response, while neuronal populations remain preserved within the tumor milieu.

### GG networks exhibit seizure susceptibility and progressive divergence of activity patterns with maturation

To gain further insight into the link between GGs and epileptic seizures, we next investigated neuronal population and seizure-like activities across groups and in a spatially dependent manner using SpikeSpector analytical tool (Cases-Cunillera et al., 2026). For this purpose, we performed multielectrode array (MEA) recordings on brain slices from GG and Ctrl mice at p5 and p20. First, we assessed the occurrence of interictal discharges (IIDs), during basal aCSF perfusion. IIDs were detected in GG^p5^ and GG^p20^ tissues, but also in Ctrl tissues (**Fig. 2A**). We next quantified the average number of IIDs per electrode (during 500 s recordings) and the proportion of active electrodes per slice (n=3-7, from 3-5 mice). Although GG^p5^ tissues tended to exhibit a higher number of IIDs per electrode and a larger fraction of active electrodes compared with Ctrl^p5^ tissues, these differences did not reach statistical significance. Similarly, no significant differences were observed between GG^p20^ and Ctrl^p20^ conditions for either metric. These results indicate that IID events are present in both control and GG tissues, with comparable levels of overall event occurrence and network participation (**Fig. 2B**). To determine whether the electrophysiological characteristics of these population events differed despite similar overall occurrence rates, we next analyzed the distributions of IID features across groups and developmental stages. Density histograms revealed that IIDs recorded from GG tissues exhibited shorter durations and half-widths, together with steeper slopes and larger amplitudes, than those recorded from control tissues at both p5 and p20 (**Fig. 2C**). These findings indicate that GG tissues generate interictal-like discharges that are faster and larger in magnitude, consistent with increased neuronal synchrony and network hyperexcitability, two hallmarks of epileptogenic circuits.

**Figure 2.**
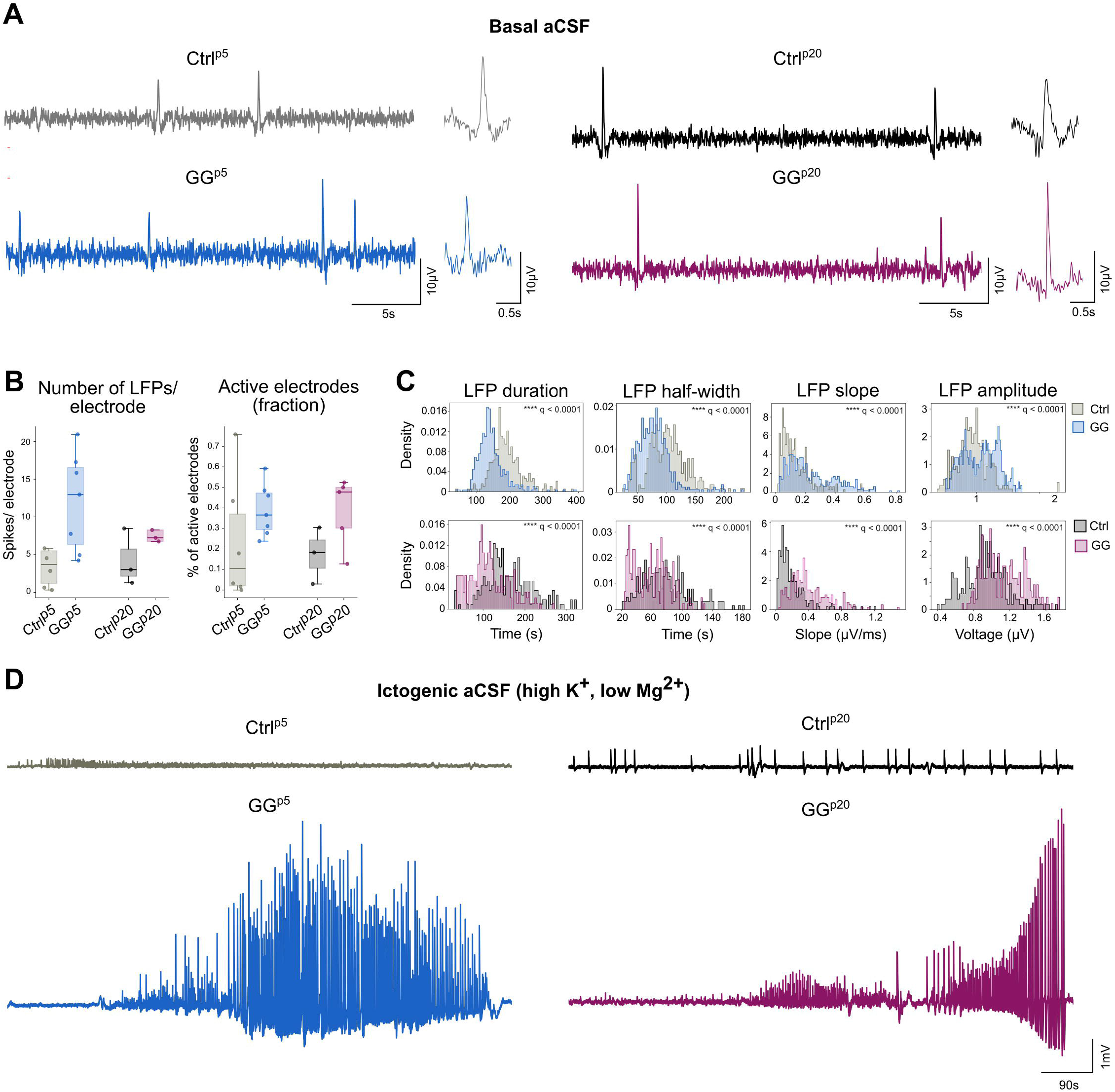
Electrophysiological characterization of IID and ID activity in Ctrl and GG slices at p5 and p20. (**A**) Representative voltage traces in basal aCSF showing IIDs in Ctrl^p5^, GG^p5^, Ctrl^p20^, and GG^p20^ slices. (**B**) Quantification of the number of IIDs per electrode and fraction of active electrodes during IID activity in aCSF conditions (n=3-7 slices, from 3-5 mice), Welch’s two-sided t-tests compared Ctrl vs GG within each age. (**C**) Density histograms showing the distribution of IID duration, half-width, slope, and amplitude across groups at p5 (top) and p20 (bottom). GG slices displayed altered distributions of IIDs features compared to Ctrl tissues (n=5-6, from 3-5 mice). Statistical significance between distributions was assessed using Kolmogorov– Smirnov tests followed by false discovery rate (FDR) correction for multiple comparisons (**** q < 0.0001). (**D**) Representative traces under pro-epileptiform conditions (ictogenic aCSF with high K^+^, low Mg^2+^) illustrating ictal-like discharges (ID).

Finally, we sought to assess the epileptic vs physiological nature of the population events by treating the brain slices with an ictogenic solution (aCSF: high K^+^, low Mg^2+^) in order to elicit seizure-like events. Strikingly, seizure-like events were significantly more frequent in GG, compared to Ctrl, brain slices at both p5 (8 out of 8 for GG, 1 out of 6 for Ctrl), and p20 (8 out of 8 for GG, 1 out of 7 for Ctrl, **Fig. 2D**), validating that BRAF^V600E^/Akt^A^ tumors can trigger seizure-like activies at very early stages of development that persist into young ages.

### Developmental shift of seizure initiation toward tumor-proximal regions in GGs

Next, we aim to investigate the site of origin of seizure activity in the GG brain slices, taking into account IUE/tumor- and GFAP/astrocytic-positive areas to distinguish tumoral vs peritumoral areas. For the region around each electrode (0.04mm²), we quantified the percentage of stained area for mCherry (marker of IU-electroporated cells) and GFAP (marker of astrocytes) in both GG^p5^ (**Fig. 3A**) and GG^p20^ (**Fig. 3C**). Continuous MEA recordings captured the transition from IIDs to preictal discharges (PIDs) and finally ictal discharges for the seizure onset (ID-SO), allowing us to examine how the spatial distribution of epileptiform activity evolved throughout seizure generation. Our data showed that in GG^p5^ slices, PIDs and ID-SO typically originated in peritumoral brain regions with low mCherry and high GFAP staining, and then propagated into both the adjacent cortex and the tumor itself (**Fig. 3B**). In contrast, for the GG^p20^ slices, PIDs and ID-SO occurred in brain regions more adjacent to the tumor tissue and they propagated to the neighbouring cortex (**Fig. 3D**).

**Figure 3.**
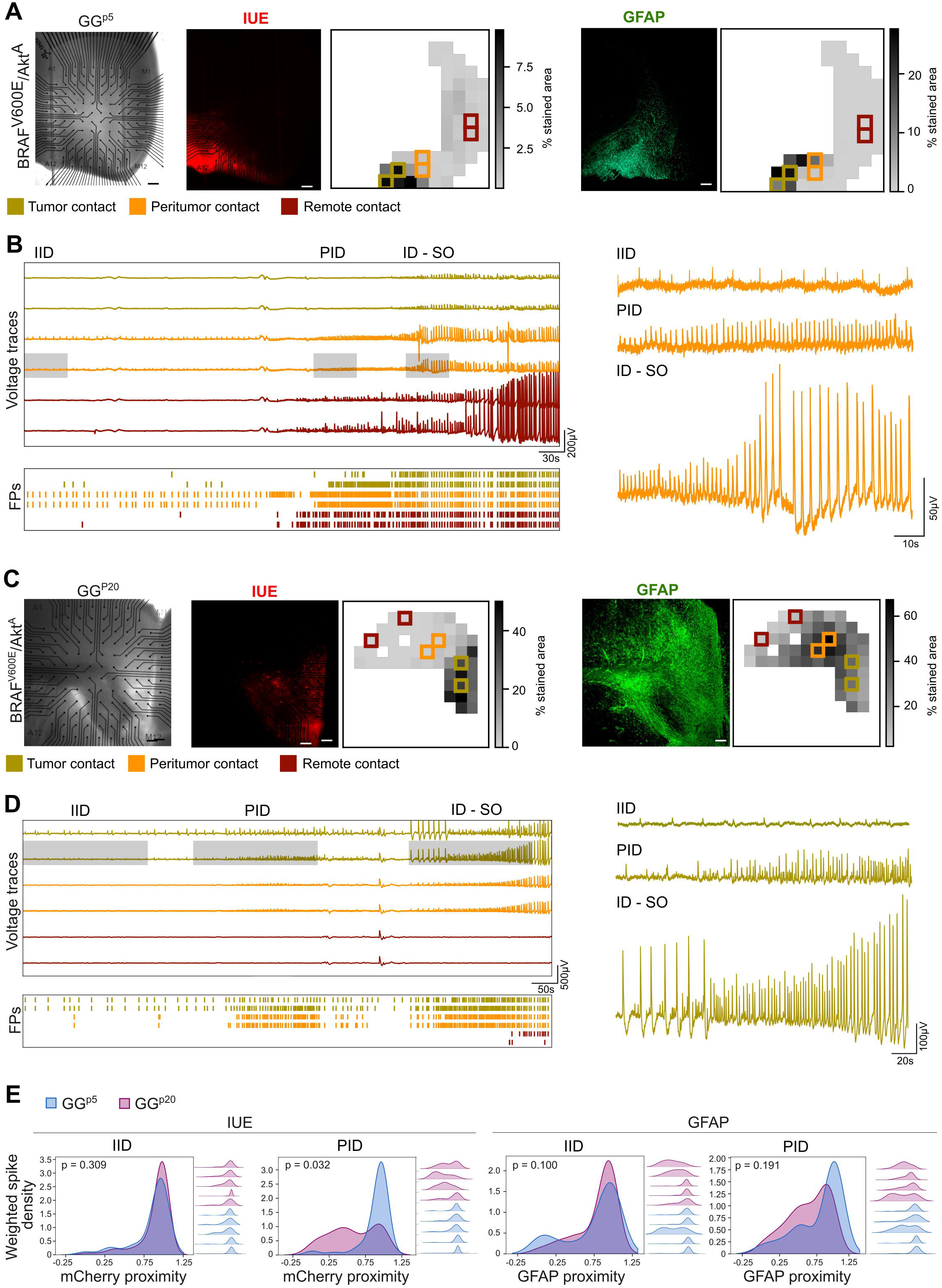
Spatial mapping of spike activity relative to IUE-labeled and GFAP-positive regions in GG^p5^ and GG^p20^. (**A**, **C**) Representative images from GG^p5^ (**A**) and GG^p20^ (**C**) slices expressing BRAF^V600E^/Akt^A^. Left panels show MEA electrode layout overlaid on brightfield images. IUE (mCherry, red) and GFAP (green) fluorescence images are shown together with corresponding spatial maps of the percentage of stained area for each 0.04m^2^ square. Colored squares indicate electrodes selected for representative traces. Scale bar, 200μm). (**B**, **D**) Representative voltage traces recorded with MEA from selected electrodes in GG^p5^ (**B**) and GG^p20^ (**D**), illustrating IID, PID, and ID - seizure onset (SO) event patterns (highlighted shaded gray rectangles). Bottom panels show raster plots of detected events over time. Traces are color-coded according to electrode location relative to tumor regions: dark red tones indicate electrodes far from the IUE-positive area (remote contact), orange tones indicate electrodes at the peritumoral border (peritumoral contact), and olive green tones indicate electrodes within IUE-positive regions (tumor contact). Right panels display higher temporal resolution examples of PID and ID activity patterns. (**E**) Density plots show the distribution of weighted spike counts as a function of distance to mCherry signal (left panels) and GFAP signal (right panels) for IID and PID events. Right-side ridge plots illustrate the distribution of proximity values across recordings for each individual slice. Group differences between p5 and p20 were evaluated using two-sided Mann–Whitney U tests, and slice-level Spearman p-values< 0.05 were considered statistically significant.

To determine whether the relationship between spiking activity and staining changed with age, we computed, for each slice, the Spearman correlation between spike count and either mCherry/IUE or GFAP staining values for IIDs as well as PIDs. Spike counts during PID periods were used to characterize the spatial distribution of activity associated with the early ictal recruitment period preceding ID-SO. These correlations were then compared between p5 and p20 brain slices (n=4-5, from 3-4 mice). Our results show that while the occurrence of IIDs did not change between p5 and p20 with respect to the mCherry/IUE or GFAP proximity, seizure initiation differed between developmental timepoints. Analysis of the mCherry–spike count correlations revealed a significant developmental difference for PIDs and ID-SO, with higher spike counts occurring closer to the tumor in p20 compared to p5 GG brain slices. Regarding GFAP-spike correlations, although higher spike counts tended to occur closer to GFAP-positive regions at p20 compared to p5, this difference was not statistically significant for PIDs nor for ID-SO (**Fig. 3E**). Overall, these results suggest that the spatial relationship between spike activity and tumor cells changes with maturation, with SO events in GG^p20^ occurring closer to tumor cells harboring the mutations than in GG^p5^, which were rather generated by remote areas.

### Human GG brain slices reveal seizure activity in the peritumoral cortex without histopathological alterations

We next aim to translate our previous results on human cortical brain slices from GG patients (6 slices from 3 patients). We restricted this analysis to assess whether the cortical tissue adjacent to GG was epileptic. For this purpose, we used MEA to record from human brain slices, which were subsequently stained with GFAP in order to locate the astrocytic lesion (**Fig. 4A** and **C**).

**Figure 4.**
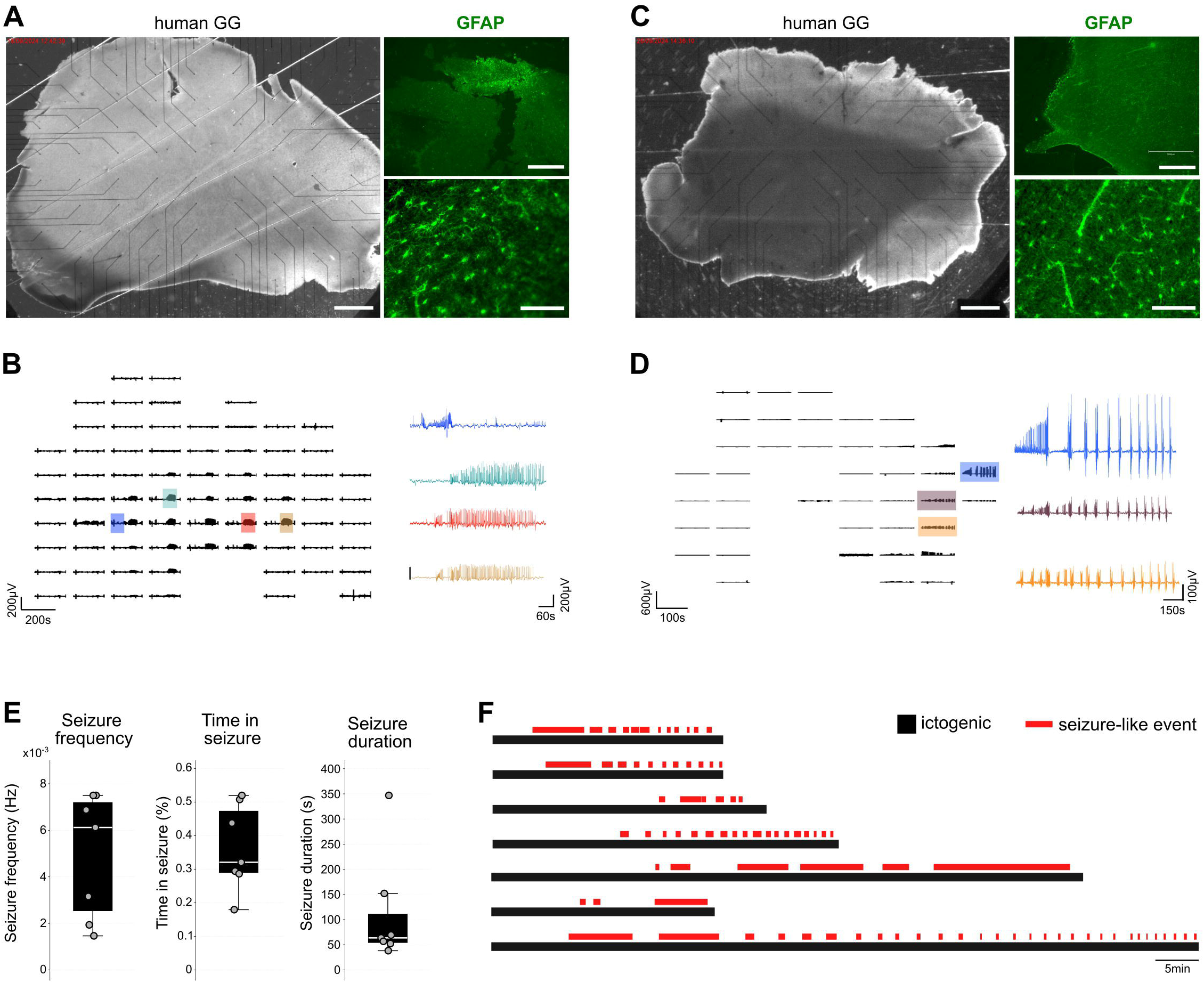
Representative electrophysiological recordings from human GG brain slices. (**A**, **C**) Representative brightfield images of acute human GG brain slices from two independent patients (scale bar, 1500μm), with corresponding GFAP immunofluorescence images shown at low (scale bar, 1250μm) and higher (scale bar, 150μm) magnifications. (**B**, **D**) Representative MEA recordings from the respective GG slices. Left panels show multi-channel voltage activity over time. Colored insets indicate selected recording regions, and right panels display higher temporal resolution traces from the indicated channels. (**E**) Quantification of seizure-like event properties, including seizure frequency (Hz), percentage of time spent in seizure activity, and seizure duration (s). Individual data points represent independent slice recordings. (**F**) Temporal distribution of MEA recordings under ictogenic solution (black) and seizure-like events (red) across recorded human GG slices.

The seizing slices were assessed in a spatially dependent manner, and our results showed that seizure activity was present in peritumoral regions (**Fig. 4B** and **D**). These results suggest that GGs can alter neuronal network activity beyond the tumor core, extending into adjacent cortical regions. This is consistent with our mouse GG data, where we observe that the neighboring cortex is capable of propagating seizure activity at both p5 and p20 (**Fig. 3B** and **D**). Quantification showed variable seizure frequency, time spent in seizure, and seizure duration across recorded slices, reflecting heterogeneous epileptiform activity among GG samples (**Fig. 4E**). Raster-style event mapping further demonstrated repeated seizure-like events occurring during ictogenic recording periods, with some slices showing prolonged and recurrent epileptiform episodes (**Fig. 4F**).

### Developmental arrest of intrinsic electrophysiological maturation in tumor neurons and progressive hyperexcitability of the peritumoral network

To assess how GGs impact neuronal intrinsic properties and local network excitability, we performed whole-cell patch-clamp recordings from IUE-positive tumor (GG+) or control (Ctrl+) neurons and from IUE-negative neurons adjacent to the IUE region (referred to as GG- or Ctrl-) in both p5 and p20 brains (**Fig. 5A**). At p5, intrinsic membrane properties, including resting membrane potential (RMP), membrane resistance (Rm), and membrane capacitance (Cm), were comparable across all recorded groups, indicating a shared electrophysiological state at early postnatal stages. By p20, while Ctrl+, Ctrl-, and GG-neurons exhibited the expected maturation-associated changes in intrinsic membrane properties, distinct from p5, GG+ neurons retained electrophysiological profiles resembling those observed at p5, indicating a developmental arrest of intrinsic membrane maturation. Statistical differences were observed between GG+ and Ctrl+ (positive for IUE) for the resting membrane potential and membrane resistance (**Fig. 5B**). We next examined excitatory postsynaptic potentials (ePSCs) to assess functional synaptic connectivity. At p5, control (Ctrl+ and Ctrl-) and peri-tumoral neurons (GG-) displayed spontaneous synaptic activity, whereas GG+ neurons exhibited a more silent phenotype with sparse, very small ePSCs (**Fig. 5C** and **E**). Moreover, peri-tumoral GG-neurons at p5 showed similar spontaneous synaptic activity compared to controls. By p20, peritumoral GG-neurons adjacent to the tumor exhibited strikingly elevated ePSC frequency, amplitude, and total integrated synaptic activity relative to all other groups (**Fig. 5D** and **E**), consistent with pronounced peritumoral network hyperexcitability.

**Figure 5.**
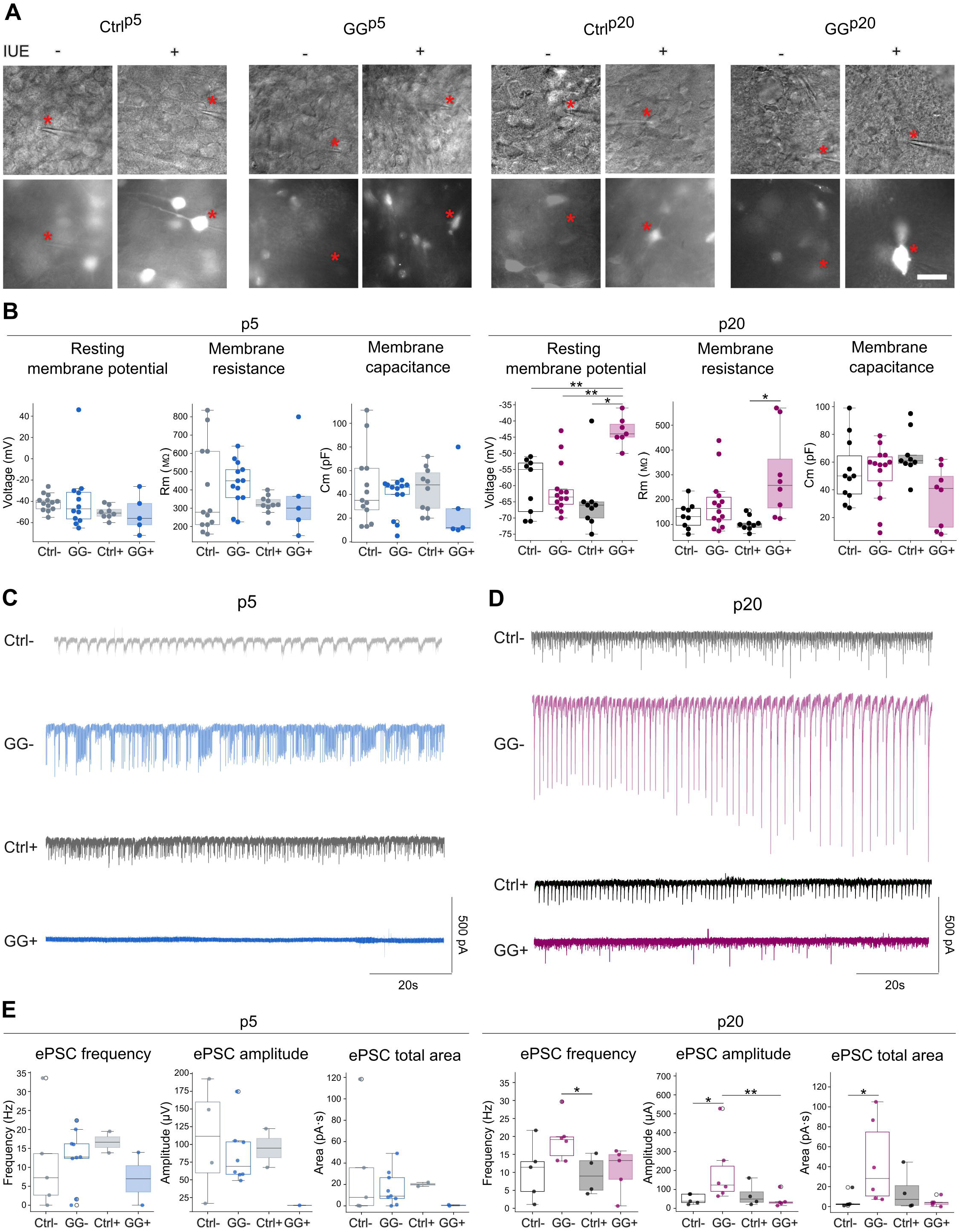
Representative electrophysiological recordings and intrinsic membrane properties of neurons in Ctrl and GG tissue. (**A**) Representative images of recorded cells in Ctrl and GG cortices at p5 and p20, stratified by IUE status (IUE− and IUE+). Upper panels show brightfield images; lower panels show fluorescence identifying IUE-positive cells. Red asterisks indicate recorded neurons. Scale bar: 20μm. (**B**) Quantification of intrinsic membrane properties (resting membrane potential (RMP), membrane resistance (Rm) and membrane capacitance (Cm) at p5 (left) and p20 (right), in Ctrl and GG neurons, stratified by IUE status. Each dot represents one recorded cell (n = 5-13, from 3-5 mice). Pairwise comparisons were performed using two-sided Mann–Whitney U tests with Holm correction for multiple testing. p < 0.05 (*) and p < 0.01 (**). (**C**, **D**) Representative traces of sEPSCs at p5 (**C**) and p20 (**D**) from Ctrl and GG with and without IUE. (**E**) Quantification of sEPSC recorded in neurons from Ctrl and GG cortical slices at p5 and p20. Statistical analyses were performed using Welch’s t-test for normally distributed datasets or Mann–Whitney U tests for non-parametric datasets following Shapiro–Wilk normality testing. Statistical significance is indicated as follows: * p < 0.05, ** p < 0.01.

Together, these data reveal a functional dichotomy within the GG microenvironment, in which tumor-resident neurons remain developmentally immature with limited synaptic integration, whereas adjacent neurons progressively acquire a hyperexcitable phenotype, providing a direct electrophysiological substrate for tumor-associated epileptogenesis.

### Single-nucleus RNA sequencing reveals stage-specific neuronal transcriptional remodeling in the peritumoral microenvironment

To define how GGs transcriptionally remodel their surrounding tissue across development, we performed single-nucleus RNA sequencing on peritumoral and peri-control cortical tissue at p5 and p30, rather than at p20 for technical reasons. Unsupervised clustering of all nuclei revealed clear segregation of the major brain cell populations, including excitatory neurons, inhibitory neurons, astrocytes, oligodendrocytes, OPCs, microglia and endothelial cells (**Fig. 6A**). When datasets were analyzed separately by developmental stage, clustering structure and major cell identities were highly preserved at both p5 and p20 (**Fig. 6B**), confirming robustness of cell-type annotation across time points. Relative cell-type composition, visualized by stacked bar plots, showed no major shifts in overall cellular proportions between peritumoral and peri-control samples, with the exception that astrocytes were modestly increased in GG relative to control in both time points, as expected. (**Fig. 6C**). Dot plot visualization of canonical marker genes confirmed accurate labeling of all major neural and glial populations at both time points (**Fig. 6D**).

**Figure 6.**
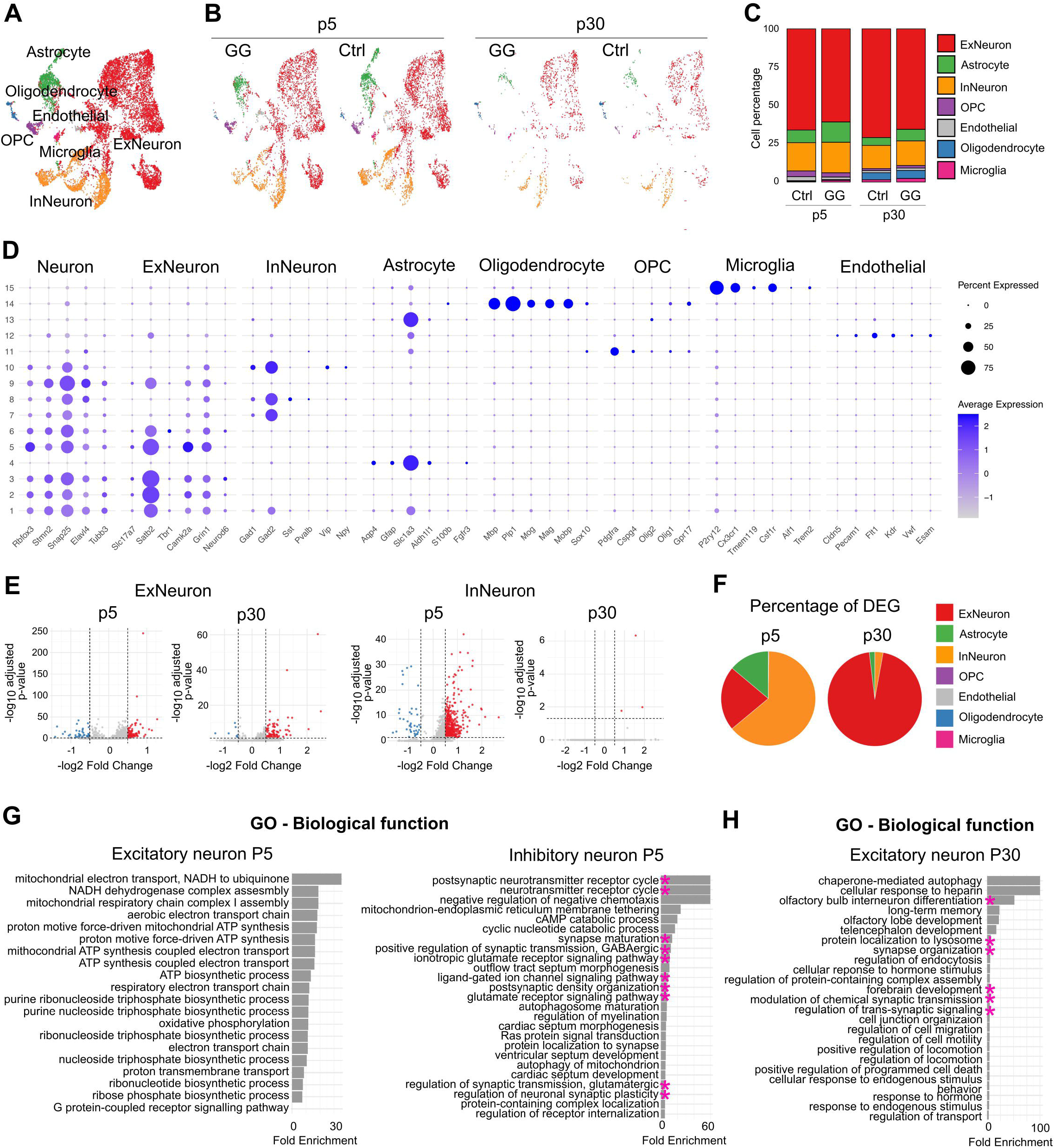
Single-nucleus RNA sequencing reveals cell-type composition and differential gene expression in GG and control cortex across developmental stages. (**A**) UMAP projection of all nuclei from Ctrl and GG samples showing clustering of major brain cell populations. (**B**) UMAP projections separated by condition (Ctrl and GG) and developmental stage (p5 and p30). (**C**) Stacked bar plots showing the relative proportions of each major cell type in Ctrl and GG samples at p5 and p30. (**D**) Dot plot showing expression of canonical marker genes used for cell-type annotation. (**E**) Volcano plots showing differential gene expression between GG and Ctrl samples for excitatory and inhibitory neuronal populations at p5 and p30. Red dots indicate significantly upregulated genes, and blue dots indicate significantly downregulated genes. Dashed lines indicate fold-change and adjusted p-value thresholds. (**F**) Pie charts summarizing the percentage of DEGs across major cell types at p5 and p30. (**G**-**H**) Gene Ontology (GO) biological process enrichment analysis of DEGs in excitatory and inhibitory neurons at p5 (**G**) and excitatory neurons at p30 (**H**). GO terms are sorted according to fold enrichment.

Differential gene expression analysis on neuronal subtypes revealed that at p5, gene expression alterations were observed in excitatory as well as inhibitory neurons (**Fig. 6E** and **F**). However, at p30, transcriptional differences were observed exclusively in excitatory neurons, while inhibitory neurons no longer displayed significant differential expression (**Fig. 6E** and **F**). Functional enrichment analysis of biological function demonstrated that at p5, only inhibitory neurons exhibited GO terms related to synaptic transmission and neuronal signaling (pink asterisks), whereas excitatory neurons displayed more limited, largely non-synaptic transcriptional changes (**Fig. 6G**). In contrast, at p30, functional GO terms associated with neuronal activity and synaptic signaling were selectively enriched in excitatory neurons (pink asterisks, **Fig. 6H**), indicating a developmental shift in tumor-associated neuronal functional remodeling from inhibitory to excitatory neuronal populations.

Together, these data demonstrate that GGs induce dynamic, developmentally staged transcriptional remodeling of peritumoral neurons, characterized by early inhibitory synaptic dysfunction and later excitatory neuronal reprogramming.

### Developmental-specific pharmacology of seizures in GG slices

Given the early functional involvement of inhibitory neuronal populations identified by transcriptomic profiling and the well-established dependence of GABAergic signaling on intracellular chloride homeostasis, we next examined whether pharmacological modulation of chloride-dependent inhibition could attenuate tumor-associated network hyperexcitability. To this end, we treated acute GG^p5^ and GG^p20^ slices with bumetanide, a selective inhibitor of the NKCC1 chloride cotransporter, under ictogenic conditions and measured extracellular activities.

In neonatal-derived GG^p5^ slices, bumetanide significantly reduced seizure frequency, duration, and total time spent in seizure compared to ictogenic baseline conditions. Washout partially restored seizure activity, indicating a reversible modulation of network excitability consistent with chloride-dependent mechanisms (**Fig. 7A**). Interestingly, young adult-derived GG^p20^ slices did not exhibit a significant response to bumetanide under identical conditions (**Fig. 7B**). Notably, bumetanide could not block seizure-like activity in slices from GG^p8-9^ mice (**Fig. 7C**), suggesting a narrow developmental window during which the treatment is effective. Together, these findings demonstrate a developmental stage–dependent sensitivity of GG-associated seizures to NKCC1 inhibition, supporting a greater contribution of chloride-dependent GABAergic signaling hyperexcitability in early postnatal tumor networks.

**Figure 7.**
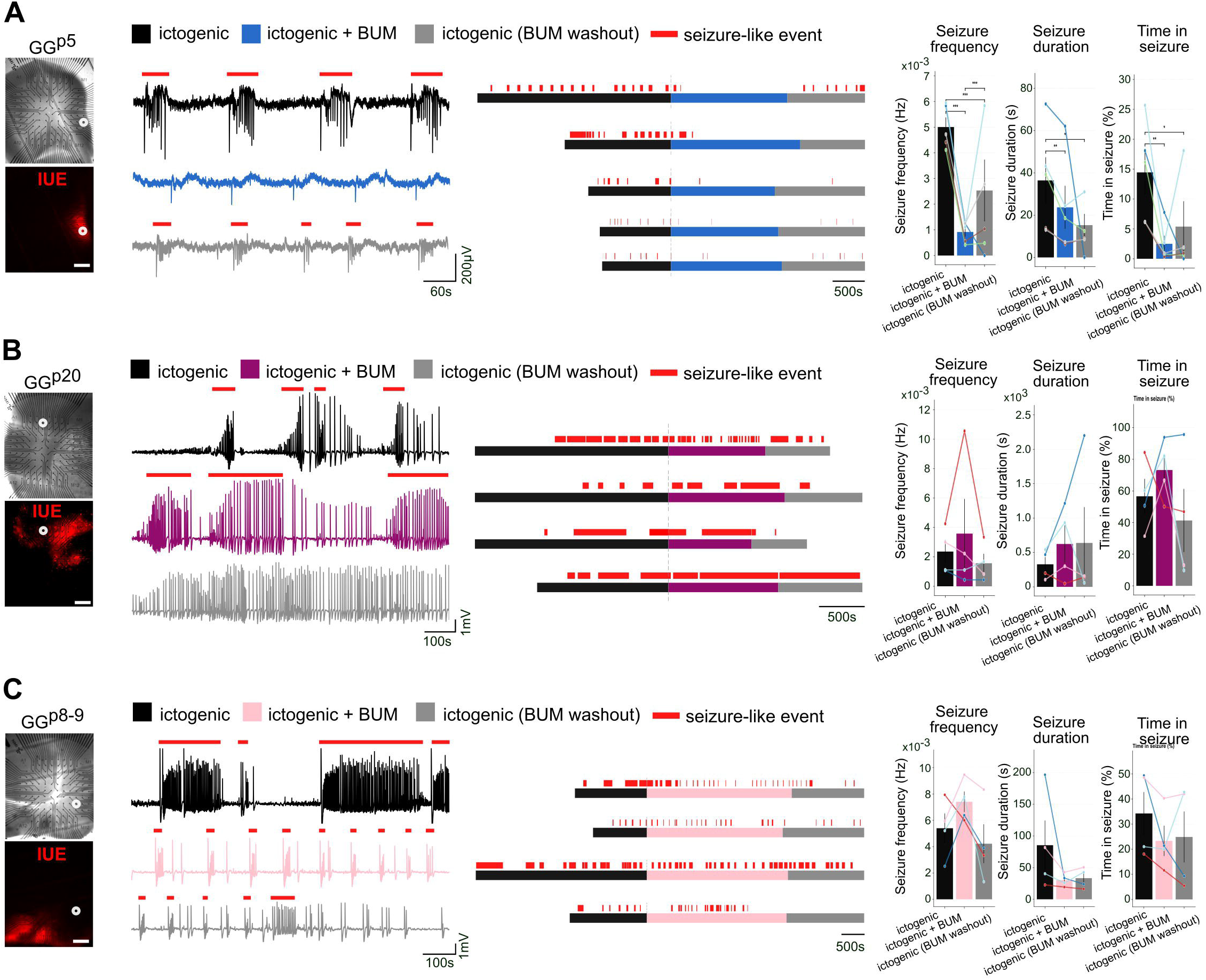
Modulation of seizure-like activity by bumetanide (BUM) in GG slices. (**A**-**C**) Representative recordings from GG^p5^ (**A**), GG^p20^ (**B**) and GG^p8-9^ (**C**) slices under ictogenic conditions (n=4-5, from 3-4 mice). Left panels show brightfield and IUE fluorescence images of recorded regions. Scale bars, 500μm. Middle panels display representative voltage traces recorded during ictogenic aCSF (black), following addition of BUM (blue) and during BUM washout (grey). Red bars indicate detected seizure-like events. Horizontal timelines summarize the duration of ictogenic exposure, BUM treatment, and washout periods. Right panels show quantification of seizure parameters, including seizure frequency, duration, and total time spent in seizure across conditions. Individual dots represent slices. Statistics: paired Friedman test. *p < 0.05, **p < 0.01, ***and p < 0.001.

## Discussion

A central finding of our study is the developmental shift in seizure initiation relative to the tumoral cells. In early postnatal slices, seizure initiation preferentially occurred in regions remote from GG cells, whereas in more mature slices, initiation shifted toward tumor-adjacent areas. This dynamic reorganization indicates that tumor–network coupling evolves with brain maturation and reflects age-dependent mechanisms of epileptogenic integration, consistent with other developmental models of pediatric epileptogenesis (Potthoff et al., 2025; Slegers and Blumcke, 2020). The early vulnerability of postnatal networks likely reflects immature inhibitory signaling and heightened susceptibility of developing neurons to tumor-induced modulation (Müller et al., 2024), whereas later seizure generation appears increasingly driven by excitatory remodeling, structural tumor–neuron integration, and inflammatory signaling. Notably, seizure initiation at P5 preferentially occurred within peritumoral regions, which were characterized by robust astrocytic reactivity already at this early developmental stage. The presence of extensive GFAP-positive astrocytes postnatally suggests that gliosis in GG is initiated early during development rather than arising solely as a consequence of chronic seizures. This response may reflect direct effects of embryonic BRAF^V600E/^Akt^A^ expression on the developing glial environment and/or the release of inflammatory mediators (Aronica and Crino, 2011; Cases-Cunillera et al., 2022; Vezzani et al., 2011). Together, these findings raise the possibility that early gliosis contributes to shaping an epileptogenic niche before the later shift toward tumor-proximal seizure onset.

At both developmental stages, seizure activity was not confined to the tumor core but involved adjacent cortex. Our spatial MEA recordings and human GG slice data demonstrate that epileptiform activity appears in peritumoral regions without tumor cell labeling, supporting the concept of a distributed epileptogenic network. This concept is consistent with neuropathological studies of LEATs showing that epileptogenicity may involve not only the tumor itself but also the surrounding cortex, which can display cytoarchitectural, inflammatory, or synaptic alterations (Aronica and Crino, 2014; Slegers and Blumcke, 2020; Thom et al., 2012). This aligns with surgical evidence showing that seizure outcome depends not only on tumor resection but also on removal of adjacent epileptogenic cortex (Blumcke et al., 2014; Giulioni et al., 2014; Thom et al., 2012), and with outcome data demonstrating persistent seizures in a subset of patients despite gross total resection. Similar tumor–network coupling and peritumoral hyperexcitability have been described in gliomas (Huberfeld et al., 2011; Pallud et al., 2014), which, in addition, are characterized by tumor–neuron synaptic interactions and circuit hyperactivity (Venkatesh et al., 2019). Together, these findings support the view that GG-associated epilepsy reflects network-level remodeling rather than a purely focal lesion.

At the cellular level, our whole-cell recordings reveal a developmental arrest of intrinsic maturation in tumor-resident neurons harboring BRAF^V600E^/Akt^A^ expression, which at p20 retained immature electrophysiological characteristics similar to p5. This observation is consistent with evidence that BRAF^V600E^ expression in neural progenitors disrupts pyramidal neuron electrophysiological properties (Goz et al., 2020; Koh et al., 2018). Our data extend these findings by demonstrating that tumor-resident neurons fail to undergo normal postnatal maturation, potentially maintaining a hyperexcitable, developmentally immature phenotype that contributes to early network instability.

Our transcriptomic analyses further support a stage-specific trajectory of tumor-associated remodeling. Studies on GG from adult mouse and human brain tissue have reported expression alterations of GABA receptor subunits and related inhibitory signaling components (Aronica et al., 2007a; Aronica and Crino, 2014; Kyriazi et al., 2023; Samadani et al., 2007), implicating impaired inhibition in tumor-associated epileptogenicity. However, these studies arise from resected brain tissue that may have been affected by prolonged epilepsy. In contrast, our developmental model reveals that early postnatal stages are characterized by prominent differential expression and functional enrichment of synaptic programs in inhibitory neurons, whereas later remodeling shifts toward excitatory neuronal and activity-related pathways. Together with our physiological data, this suggests that early inhibitory dysfunction may represent a critical window that precedes and shapes later transcriptomic signatures observed in human specimens.

Consistent with this developmental framework, bumetanide significantly reduced seizure frequency, duration, and ictal burden in neonatal GG slices but was ineffective at later stages. Given the developmental regulation of chloride transporters and GABAergic maturation, early networks are particularly dependent on NKCC1-mediated chloride gradients. Prior evidence linking disrupted chloride homeostasis to epileptogenesis in developmental tumors and dysplastic cortex (Aronica et al., 2007b) supports this interpretation. Consistent with this, pharmacological inhibition of NKCC1 with bumetanide has been shown to suppress seizure activity and epileptiform synchronization in neonatal hippocampal and other postnatal epileptogenic models (Dzhala et al., 2008). The lack of efficacy at later stages suggests that seizure mechanisms progressively shift toward excitatory synaptic remodeling and inflammatory signaling (Cases-Cunillera et al., 2022; Vezzani et al., 2011), rendering chloride-targeted intervention less effective, consistent with a maturation of GABAergic signaling and with the prominent glutamatergic mechanisms then at play. Clinically, seizure freedom after GG resection is common but not universal (Blumcke et al., 2014), supporting the notion that epileptogenic mechanisms may evolve beyond the primary lesion and become embedded within broader cortical networks. These age-dependent clinical trajectories parallel our experimental findings of stage-specific epileptogenic mechanisms and pharmacological responsiveness.

Together, our data support a unified model in which embryonically initiated tumor formation disrupts immature cortical networks, establishing an early chloride-sensitive epileptogenic state. With maturation, tumor-host interactions evolve, intrinsic maturation of tumor neurons remains arrested, transcriptional programs shift from inhibitory to excitatory remodeling, and seizure generation becomes increasingly embedded within reorganized cortical networks. Recognizing this developmental evolution may inform surgical strategy, prognostic assessment, and the design of age-specific adjunctive therapies in GG-associated epilepsy.

## Materials and methods

### Sex as a biological variable

Both male and female animals/patients were included in this study.

### Immunohistochemistry

Samples were washed three times for 5 min each with phosphate-buffered saline (PBS). Tissue permeabilization was performed by incubating the samples in PBT (PBS containing 0.2% Triton X-100) for three washes of 5 min each. Blocking was carried out using PGT solution (PBS containing 0.25% Triton X-100 and 0.2% gelatin). Samples were then incubated with the primary antibody diluted in PGT overnight at 4 °C. Following primary antibody incubation, samples were washed with PBT and subsequently incubated in PGT for 10 min at room temperature. Secondary antibodies together with DAPI were then applied in PGT and incubated for 45 minutes at room temperature in the dark. Finally, samples were washed with PBS to remove excess antibodies and mounted using an appropriate mounting medium for fluorescence imaging.

### Intraventricular in-utero electroporation

The plasmids expressing the genes used for in-utero electroporation (IUE) were developed as previously described (Cases-Cunillera et al., 2022). IUE was performed on timed-pregnant CD1 mice (Janvier Labs) at embryonic day 14.5 (E14.5). Mice were anesthetized with isoflurane (3% for induction, 2% for maintenance). A small incision was made in the abdominal wall to expose the uterine horns, which were kept hydrated with sterile PBS at 37°C throughout the procedure. A solution containing DNA plasmids (1.5 μg/μl) mixed with Fast Green FCF (Sigma Aldrich) for visualization was injected into the lateral ventricles of the embryos. Electroporation was performed using a NEPA21 Super Electroporator (Nepa Gene, Chiba, Japan), delivering five 40 V electrical pulses. Afterward, the embryos were placed back into the abdominal cavity, and the muscle and skin were sutured sequentially.

### Mouse brain slice preparation

Mice subjected to IUE were sacrificed by decapitation at either postnatal day 5 (± 2 days) or day 20 (± 2 days). Following decapitation, the brains were carefully removed and immediately transferred into an ice-cold, oxygenated cutting solution containing (in mM): 1.25 NaH_2_PO_4_, 2.5 KCl, 3.1 sodium pyruvate, 7 MgCl_2_, 10 D-Glucose, 26 NaHCO_3_, 11.6 sodium L-ascorbate, 110 Choline chloride and 0.5 CaCl_2_. Brain slices of 400 µm thickness were prepared using a vibratome and then transferred to an interface chamber, which was maintained at a temperature of 32°C, equilibrated with 95% O₂ and 5% CO₂. The chamber was perfused with an oxygenated recording solution (referred to as aCSF) containing (in mM): 3 KCl, 11.1 D-Glucose, 26 NaHCO_3_, 124 NaCl, 1.3 MgCl_2_ and 1.6 CaCl_2_. The brain slices were incubated in this interface chamber at 2 ml/min for a minimum of 60 minutes before electrophysiological recordings.

### Human brain slice preparation

Preparation of human brain slices was performed similarly to that described before (Dossi et al., 2014). Human GG specimens were immediately placed in the operating room in an iced cutting solution, equilibrated with 5% CO2 in 95% O2, and transported to the laboratory. Blood clots, meninges and vessels were carefully removed and 400 μm-thick slices were cut with a vibratome. Slices were placed in an interface chamber at 37°C and continuously perfused with aCSF at 2 ml/min for at least 1 hour.

### MultiElectrode Array (MEA) recording and data analysis

Mouse brain slices were transferred to the recording chamber containing an array of 120 planar titanium nitride-coated electrodes (Multi Channel Systems). Each electrode measures 30 µm in diameter, with 200 µm spacing between them. The brain slices, containing both tumor and peritumoral regions, were precisely positioned on the electrode grid by visualization of the fluorescence signal emitted from tumor cells using a Fluorescent Probe in Flashlight Set (Laborimpex). Human brain slices were recorded on a bigger recording chamber containing an array of 120 electrodes with an electrode diameter of 30 µm, and an inter-electrode interval of 1500 µm / 1000 µm. A custom-made harp was used to ensure tight contact between the brain slice and the MEA electrodes and slices were perfused with pre-warmed (37 °C) oxygenated aCSF or ictogenic solution (aCSF with 0mM Mg^2+^ and 8mM K^+^) at a rate of 6 ml/min. For the bumetanide treatment, 8 μM bumetanide (Sigma-Aldrich B3023) was diluted in ictogenic solution during MEA recording.

Signals were recorded at a sampling rate of 10 kHz, using a MEA2100-120 system (Multichannel Systems). Commercially available software MC Rack (Multichannel Systems) was employed for data acquisition. Subsequent data analysis was performed with the SpikeSpector software analytical tool (Cases-Cunillera et al., 2026). Raw MEA signals were filtered using a low-pass filter at 25 Hz for visualization of population field events and at 1 Hz for quantitative analysis. IID and seizure-like events were detected semi-automatically based on their characteristic waveform morphology, consisting of transient field potential deflections with a well-defined onset, peak, and return to baseline. All automatically detected events were subsequently reviewed to exclude artifacts and ensure accurate event classification. For each validated event, duration, half-width, slope, and peak amplitude were extracted for further analysis.

### Patch-clamp recordings

Whole-cell patch-clamp recordings were performed using a Zeiss AxioExaminer microscope equipped with a 40× high-aperture objective and motorized micromanipulators (Luigs & Neumann). Acute brain slices were continuously superfused with carbogenated aCSF maintained at 37°C throughout the recordings. Target neurons were visually identified using infrared differential interference contrast (IR-DIC) optics and selected based on [selection criteria]. Whole-cell current-clamp and voltage-clamp recordings were obtained using borosilicate glass electrodes (3–5 MΩ) filled with an intracellular solution containing (in mM): K-gluconate 120, KCl 20, MgCl₂ 2, EGTA 0.6, MgATP 2, NaGTP 0.3, HEPES 10, and phosphocreatine 7. Recordings were acquired using a MultiClamp 700A amplifier (Axon Instruments, Union City, CA, USA). Signals were filtered at 4 kHz and digitized using pClamp 10 software (Molecular Devices, Sunnyvale, CA, USA). Voltage-clamp recordings were performed at a holding potential of −70 mV unless otherwise stated. Series resistance was continuously monitored by applying −10 mV voltage steps throughout the experiment, and recordings were excluded if the series resistance changed by threshold, e.g., >20%.

Following establishment of the whole-cell configuration, resting membrane potential was measured, and passive membrane properties, including access resistance, membrane resistance, and membrane capacitance, were determined using the Membrane Test function in pClamp 10. Spontaneous synaptic events were then recorded in voltage-clamp mode. Electrophysiological data were analyzed using Clampfit (Molecular Devices) and custom Python scripts.

### Single-nuclei RNA sequencing

Single-nucleus RNA sequencing (snRNA-seq) libraries were prepared using the 10x Genomics Chromium NextGEM Single Cell 3′ Reagent Kit v3.1 according to the supplier’s protocol. Nuclei were loaded onto the Chromium GEM Chip at a concentration of 500-1,500 nuclei/µL, with a target capture of approximately 16,500 nuclei per sample. Sequencing was conducted on an Illumina NovaSeq 6000 instrument using a 28/10/10/90 bp read configuration, resulting in a mean sequencing depth of about 25,000 reads per nucleus.

### Single-nucleus RNA data processing

Raw sequencing data were processed using Cell Ranger (v9.0.1, 10x Genomics). FASTQ files were aligned to the mouse reference genome (mm10; 10x Genomics reference package refdata-gex-mm10-2024-A) using the cellranger count pipeline with default parameters. This workflow included read alignment, filtering, cell barcode processing, and unique molecular identifier (UMI) quantification to generate gene-by-cell count matrices. Downstream analyses were performed in R using Seurat (v4). Cells with fewer than 200 or more than 3,000 detected genes, or with >10% mitochondrial reads, were excluded. Data were normalized, scaled, and integrated using the standard Seurat workflow to correct for batch effects. Principal component analysis (PCA) was performed, followed by shared nearest-neighbor graph construction and Louvain clustering. Clusters were visualized using UMAP and annotated based on canonical marker genes. Differentially expressed genes were identified using the Wilcoxon rank-sum test (min.pct = 0.1) and analyzed for Gene Ontology enrichment using the PANTHER tool.

### Statistics

Statistical analyses were performed using Python. Data normality was assessed using the Shapiro–Wilk test. Comparisons between two groups were performed using Welch’s t-test or Mann–Whitney U test, as appropriate. IIDs and PIDs feature distributions were compared using Kolmogorov–Smirnov tests followed by Benjamini–Hochberg FDR correction. Seizure incidence was compared using Fisher’s exact test. Correlations between spike counts and staining intensity were assessed using Spearman’s rank correlation. Repeated-measures pharmacological experiments were analyzed using the Friedman test. Differential gene expression analysis was performed using the Wilcoxon rank-sum test with Benjamini–Hochberg correction. Statistical significance was defined as p < 0.05.

### Study approval

Human GG tissue was obtained from operations on patients from 2 adult and 1 neurosurgical centers (GHU Paris Psychiatrie et Neurosciences - Sainte-Anne, Rothschild Foundation and Necker hospitals, Paris). All patients gave written consent, and our protocol was approved by the Comité d’Evaluation et d’Ethique de l’INSERM (IRB00003888 – Protocole Neurotissus 21-864). Animal experiments were conducted at the Institut de Psychiatrie et Neurosciences de Paris (INSERM U1266), an authorized animal user establishment (authorization No. D 75-14-03), in accordance with the European Directive 2010/63/EU and French regulations on the protection of animals used for scientific purposes.

## Data availability

The data that support the findings of this study are available from the corresponding author upon request. The gene expression data have been deposited in the Gene Expression Omnibus (GEO) under accession number GSE338504.

## Acknowledgments

We gratefully acknowledge the invaluable technical assistance provided by the INSERM staff.

## Conflict of interest statement

The authors have declared that no conflict of interest exists.

## Funding

This work was supported in part by the European Research Council to GH (Consolidator grant #865592). The work related to the single-nuclei RNA sequencing was supported by the Deutsche Forschungsgemeinschaft (DFG) Research Infrastructure West German Genome Center (project 407493903) as part of the Next Generation Sequencing Competence Network (project 423957469). Next Generation Sequencing analyses were carried out at the production site Bonn (West German Genome Center Bonn/Next Generation Sequencing Core Facility of the Medical Faculty at the University of Bonn). Albert Becker received the Deutsche Forschungsgemeinschaft (DFG) grant BE 2078/8-1 as part of the DFG Sequencing call SEQ1196. The first author gratefully acknowledges support from Deutsche Krebshilfe (case number: 70115733), which funded the own postdoctoral research position associated with this work.

## Author contributions

SCC, LD, JPa, TB, RS, ED, AJB, and GH conceptualized and designed the study. SCC, LD, JR, BDF, JPy, JN, KS, AE, DF, JPa, TB, MLVQ, ED, and GH developed the methodology and conducted experiments. SCC, LD, JR, BDF, KS, and AE performed data analysis and validation. SCC, LD, and GH acquired data, curated the data and generated figures and visualizations. SCC and GH wrote the original manuscript draft. SCC, BDF, RS, ED, AJB, and GH contributed to manuscript review and editing. RS, MLVQ, ED, AJB, and GH supervised the study. GH acquired funding and administered the project. All authors reviewed, edited, and approved the final version of the manuscript.

## Notes

### Competing Interest Statement

The authors have declared no competing interest.

